# Dominance rank-associated immune gene expression is widespread, sex-specific, and a precursor to high social status in wild male baboons

**DOI:** 10.1101/366021

**Authors:** Amanda J. Lea, Mercy Y. Akinyi, Ruth Nyakundi, Peter Mareri, Fred Nyundo, Thomas Kariuki, Susan C. Alberts, Elizabeth A. Archie, Jenny Tung

**Affiliations:** Department of Biology, Duke University, Box 90338, Durham, NC 27708, USA; Institute of Primate Research, National Museums of Kenya, P. O. Box 24481, Karen 00502, Nairobi, Kenya; Department of Evolutionary Anthropology, Box 90383, Durham, NC 27708, USA; Department of Biological Sciences, University of Notre Dame, Notre Dame, Indiana 46556, USA; Duke University Population Research Institute, Box 90420, Durham, NC 27708, USA

**Keywords:** dominance hierarchy, social behavior, sociogenomics, expression quantitative trait loci, lipopolysaccharide, Mendelian randomization

## Abstract

In humans and other hierarchical species, social status is tightly linked to variation in health and fitness-related traits. Experimental manipulations of social status in female rhesus macaques suggest that this relationship is partially explained by status effects on immune gene regulation. However, social hierarchies are established and maintained in different ways across species: while some are based on kin-directed nepotism, others emerge from direct physical competition. We investigated how this variation influences the relationship between social status and immune gene regulation in wild baboons, where hierarchies in males are based on fighting ability but female hierarchies are nepotistic. We measured rank-related variation in gene expression levels in adult baboons of both sexes at baseline and in response to *ex vivo* stimulation with the bacterial endotoxin lipopolysaccharide (LPS). We identified >2000 rank-associated genes in males, an order of magnitude more than in females. In males, high status predicted increased expression of genes involved in innate immunity and preferential activation of the NFkB-mediated pro-inflammatory pathway, a pattern previously associated with low status in female rhesus macaques. Using Mendelian randomization, we reconcile these observations by demonstrating that high status-associated gene expression patterns are precursors, not consequences, of high social status in males, in support of the idea that physiological condition determines who attains high rank. Together, our work provides the first test of the relationship between social status and immune gene regulation in wild primates. It also emphasizes the importance of social context in shaping the relationship between social status and immune function.

**SIGNIFICANCE:** Social status predicts fitness outcomes in social animals, motivating efforts to understand its physiological causes and consequences. We investigated the relationship between social status and immune gene expression in wild baboons, where female status is determined by kinship but male status is determined by fighting ability. We uncover pervasive status-gene expression associations in males, but not females. High status males exhibit high levels of pro-inflammatory gene expression, in contrast to previous findings in hierarchies that are not competitively determined. Using Mendelian randomization, we show that this status-associated variation precedes dominance rank attainment: males who compete successfully for high status are already immunologically distinct. The nature of social hierarchies thus fundamentally shapes the relationship between social status and immune function.

## INTRODUCTION

Many mammalian societies are characterized by strict dominance hierarchies, in which consistent, asymmetric agonistic relationships exist between group members. In such species, position in the social hierarchy (i.e., dominance rank) often predicts physiological outcomes, such as steroid hormone levels and immune function, and components of fitness, including fertility and survival rates (1–4). Notably, the relationship between social status, health, and components of fitness is especially robust in humans (5). Across human populations, socioeconomic status (SES) has been consistently associated with variation in mortality rates (6–8), and expected lifespan can differ between the highest and lowest status individuals by a decade or more (6). While human SES has no exact equivalent in animals, dominance rank in social mammals is arguably the closest parallel (2, 4). Thus, studies of rank-related gradients are important not only for understanding the evolution of social behavior in general, but also for understanding the causes and consequences of social inequality in our own species.

The molecular signature of social status is of particular interest because it can help address the long-standing puzzle of how social gradients ‘get under the skin’ (9). Advances in this area have primarily come from a few systems (10): observational studies in humans, which have established consistent, correlative relationships between social status and gene regulation (11); models of social defeat in laboratory rodents, which have connected socially induced stress to specific neural and endocrine signaling pathways (12, 13); and experimental manipulations of dominance rank in captive rhesus macaques, which have demonstrated causal effects of social status on gene regulation under controlled conditions (14, 15). For example, by manipulating dominance rank in replicate social groups of female rhesus macaques, Snyder-Mackler et al. (14) demonstrated that low social status leads to constitutive up-regulation of genes involved in innate immune defense and inflammation. Low status females responded more strongly to *ex vivo* antigen stimulation and preferentially activated an NFkB-mediated pro-inflammatory response pathway in favor of an alternative, type I interferon-mediated pathway. Studies in humans have also correlated low SES and other sources of social adversity with up-regulation of pro-inflammatory genes and down-regulation of genes involved in the type I interferon response (11), suggesting that the transcriptional response to social adversity is conserved across species (16). Because chronic inflammation is a hallmark of aging and disease risk (17), this response has in turn been hypothesized to mediate social status gradients in health and disease susceptibility (16).

Such studies suggest that social status effects on health and fitness are mediated, at least in part, through changes in gene regulation in the immune system. However, the relationship between social status and gene regulation remains unexplored in natural populations, even though the behavioral, physiological, and fitness correlates of social status have been extensively investigated in such populations (2, 4, 18, 19). Addressing this gap is important for two reasons. First, with the exception of humans, the extent to which social status-gene regulation associations arise under non-experimental conditions is unknown, leaving open the question of whether and how they contribute to social status-fitness effects in the wild. Second, social hierarchies vary in how they are established, enforced, and maintained (2, 3, 18), and laboratory systems capture only a narrow spectrum of this variation. For example, female rhesus macaques—one of the most important models for studying social hierarchies in the lab (20)— exhibit some of the most inequitable social relationships in their genus (21). Such variation influences the effects of social status on HPA axis physiology (22) and is likely to influence its molecular signature as well.

Indeed, at least two distinct types of hierarchies are found in social mammals: (i) those based on individual competitive ability, where the main predictors of rank are age, strength, and/or body size (e.g., male olive baboons (23), male red deer (24), female African elephants (25)) and (ii) ‘nepotistic’ hierarchies, in which individuals acquire predictable and stable dominance ranks similar to those of their kin, typically because relatives support one another in agonistic encounters (26). Nepotistic hierarchies occur most often in female social mammals, especially species in which females do not disperse (e.g., Japanese macaques (27), yellow baboons (28), spotted hyenas (29)). However, nepotistic hierarchies and competitive ability-based hierarchies can co-exist in the same species due to either sex or population differences. For example, male yellow baboons compete for rank via direct physical competition, such that the highest ranking males are usually prime-age males in excellent body condition (23). In contrast, for female yellow baboons, rank is matrilineally ‘inherited’, such that adult females tend to rank immediately below their mother (28). Contrasting hierarchy types exist between sexes and populations in humans as well: in small-scale human foraging societies, male status is often based on hunting ability (30, 31), while in 19^th^ century Finnish and Mormon populations kinship ties strongly predicted female and male social status, respectively (32, 33).

Here, we use behavioral and genomic data from a long-term study population of wild yellow baboons in Kenya (34) to investigate the relationship between social status, gene expression, and immune function in two different types of status hierarchies: the highly dynamic, competitive ability-based hierarchy in males and the relatively stable, nepotism-based hierarchy in females. Baboons are an excellent system for studying rank effects because rank dynamics are well-characterized, and rank itself predicts a number of fitness-related traits (23, 35, 36). Further, differences in rank attainment and maintenance between the sexes provide a natural contrast between dominance hierarchy types.

We set out to address three major questions: (i) Is social status associated with predictable patterns of gene expression in wild baboons and, if so, which genes and biological processes are affected? (ii) To what degree does the social status-gene regulation relationship depend on sex-specific social hierarchies? and (iii) In the competitive ability-based hierarchy characteristic of males, is variation in gene expression a consequence of male rank, or instead, does variation in gene expression precede variation in male rank (i.e., because it is a component of male quality or condition)? To do so, we measured cell type composition, genome-wide gene expression levels, genetic variation, and cytokine levels in blood collected from 61 known individuals (n=26 adult females and 35 adult males). We used an experimental *ex vivo* approach in which paired samples of cells were incubated in either the presence or absence of lipopolysaccharide (LPS: a component of Gram-negative bacterial cell membranes; Figure 1). LPS is a powerful stimulant of the Toll-like receptor 4 (TLR4)-mediated innate immune response, which has previously been shown to be sensitive to social environmental conditions (4, 11, 14). We investigated sex-specific associations between gene expression and dominance rank, and used Mendelian randomization (MR) to test for causal connections between social status and innate immune gene expression in males (37). Together, our findings expand work on the functional genomic signature of social status to wild primates and highlight its dependency on sex and/or hierarchy type. In addition, our MR analyses illustrate one way in which gene regulatory variation can be harnessed to address long-standing questions in behavioral and evolutionary ecology: in this case, about the causal relationship between dominance rank and immune function in competitive hierarchies.

**Figure 1.**
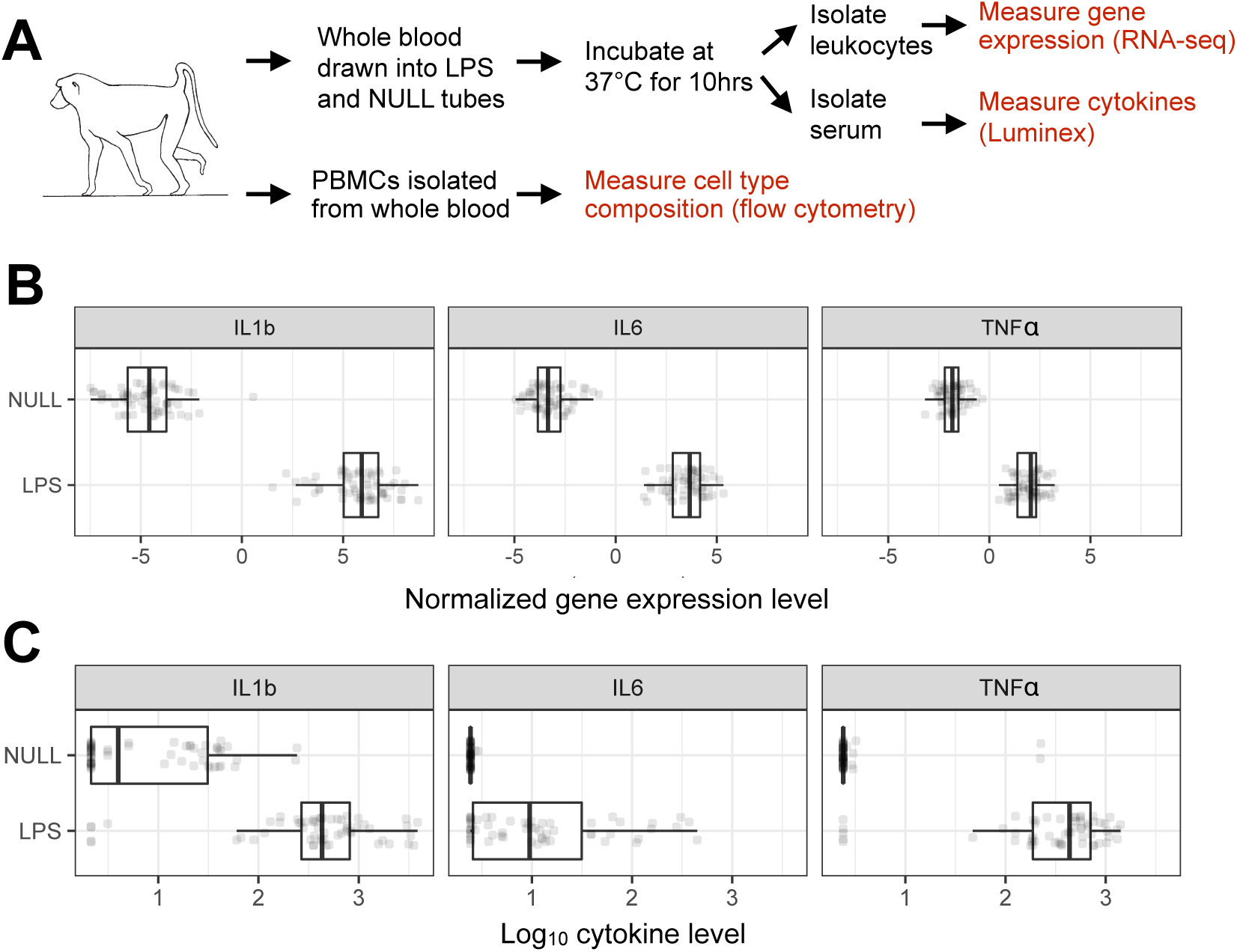
Study design. (A) Whole blood was drawn into tubes containing cell culture media alone (NULL) or media + lipopolysaccharide (LPS). Post-incubation, leukocytes were isolated for RNA-seq and serum was isolated for cytokine profiling to confirm the expected immune response. PBMCs were isolated in parallel to assess cell type composition via flow cytometry. Proinflammatory cytokines were consistently upregulated in the LPS condition at the (B) mRNA and (C) protein level (all p<10^−10^).

## RESULTS

### Ex vivo stimulation with lipopolysaccharide (LPS) induces a strong immune response

To confirm the efficacy of the LPS challenge, we measured 15 immune-related cytokines in serum collected from stimulated (LPS condition) and unstimulated (NULL condition) cells for a subset of individuals in our dataset (n=11 females, 18 males; Figure S1 and Table S1). As expected, cytokines involved in inflammatory signaling, such as IL1B (linear mixed model: p=5.41×10^−5^), IL6 (p=2.62×10^−19^), and TNFα (p=6.51×10^−11^), were strongly up-regulated in cells exposed to LPS compared to unstimulated samples from the same study subjects (Figure 1). Principal component analysis (PCA) of the full gene expression dataset (n=121 samples from 61 individuals) revealed that treatment condition was the largest source of variance in transcriptional profiles (Table S2 and Figures S2-S3). Specifically, LPS and NULL condition samples separated perfectly on PC1 (Spearman’s rank correlation, rho=0.974, p<10^−16^; Figure 2). Overall, 67% of the 7576 genes analyzed were differentially expressed between treatment conditions (linear mixed effects model, FDR<1%), and genes up-regulated in the LPS treatment condition were strongly enriched for innate immune response-related pathways (Figure S4 and Table S3; hypergeometric test, all FDR<10^−6^).

**Figure 2.**
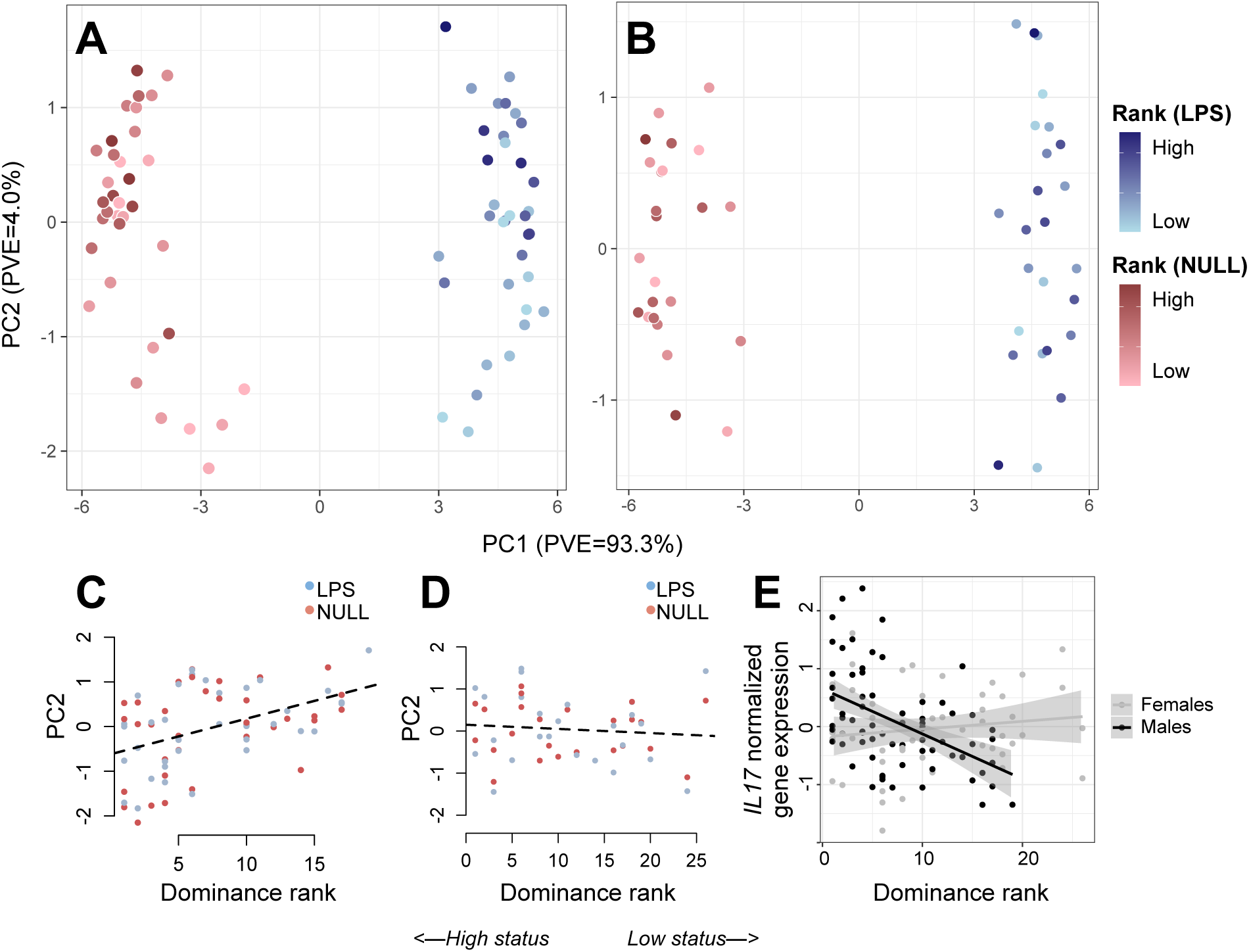
Sex-specific associations between dominance rank and gene expression. (A) PCA decomposition of LPS and NULL condition samples reveals that both treatment and dominance rank have strong effects on gene expression in males. (B) In females, treatment, but not dominance rank, affects global gene expression profiles. Legend to the right of panel B applies to both panels A and B. PC2 is correlated with rank in (C) males but not (D) females (males: rho=0.44, p=1.26×10^−4;^ females: rho=-0.11, p=0.42). (E) Example of an immune gene *(IL17)* for which rank has a strong effect in males but not females (males: rho=-0.48, p=2.13×10^−5;^ females: rho=0.12, p=0.39). In C-E, low values on the x axis signify high status and high values signify low status.

### Rank-gene expression associations are highly sex-specific

PCA including both NULL and LPS condition samples revealed a highly sex-specific signal of social status in baboon gene expression profiles. Dominance rank was significantly associated with PC2 of the gene expression data set in males (rho=0.44, p=1.26×10^−4^), but showed no such association in females, either on PC2 (rho=-0.11, p=0.42; Figure 2) or in any of the top 20 PCs (all p>0.05). At the level of individual genes, 2277 and 25 genes were associated with rank in males and females, respectively, and 1584 genes exhibited significant rank x sex interactions (FDR<5%; Figure S5 and Table S4). Sex differences in rank-gene expression associations were not primarily driven by differences in power: across 100 subsampled data sets with matched numbers of males and females, we consistently found far more rank-associated genes in males than in females (the difference in the number of genes associated with rank in males compared to females was 1387 ± 819.09 s.d.; Figure S6). Further, male and female rank effects were weakly correlated overall (Figure S5; rho=0.045, p<10^−10^), and only 4 genes were associated with rank in both sexes, no more than expected by chance (hypergeometric test: p=0.06).

Social status has been shown to influence not only steady-state gene expression levels, but also the response to immune stimulation (14, 38). Such a pattern suggests that the gene regulatory signatures of social interactions can be concealed or unmasked by other environmental factors: in female rhesus macaques, for example, dominance rank effects are magnified after LPS stimulation, because low ranking females mount stronger inflammatory responses than high ranking females (14). In contrast, in male baboons, we found relatively few genes with statistical support for a rank x treatment interaction (n=5 genes, FDR<5%) or for a rank effect on the fold change in expression levels between LPS and NULL condition cells (n=12 genes, FDR<5%). This result in part reflects low power to detect significant interactions when considering only males (Figure S7). Indeed, we observed strong evidence for male rank x condition interactions for key innate immune genes such as *IFI6* (q=0.021, p=9.57×10^−6^), *TMEM173* (q= 0.042, p=3.79×10^−4^), and *IRF9* (q= 0.047, p=7.54×10^−4^), all of which exhibited stronger responses to LPS stimulation in low ranking males relative to high ranking males. Overall, however, social status effects on gene regulation were largely consistent between LPS and NULL conditions (rho=0.619, p<10^−10^; Figure S7).

### High status predicts higher expression of innate immune genes in males

We next explored the biological function of male rank-associated genes, after filtering for genes that were up-regulated in the LPS condition (i.e., those specifically implicated in the innate immune response). Among genes up-regulated in high status males (n=478 genes; FDR<5%), we identified significant enrichment for GO terms related to the defense response, such as ‘regulation of IL6 production’ (p=2.2×10^−7^), ‘toll-like receptor signaling pathway’ (p=9.2×10^−7^), and ‘regulation of inflammatory response’ (p=2.0×10^−8^; all tests were performed relative to the background set of genes up-regulated in the LPS condition; see Figure 2, and Table S5). In contrast, genes up-regulated in low status males (n=374 genes), were enriched for GO categories related to basic cellular function and RNA processing (Figure 3 and Table S6).

**Figure 3.**
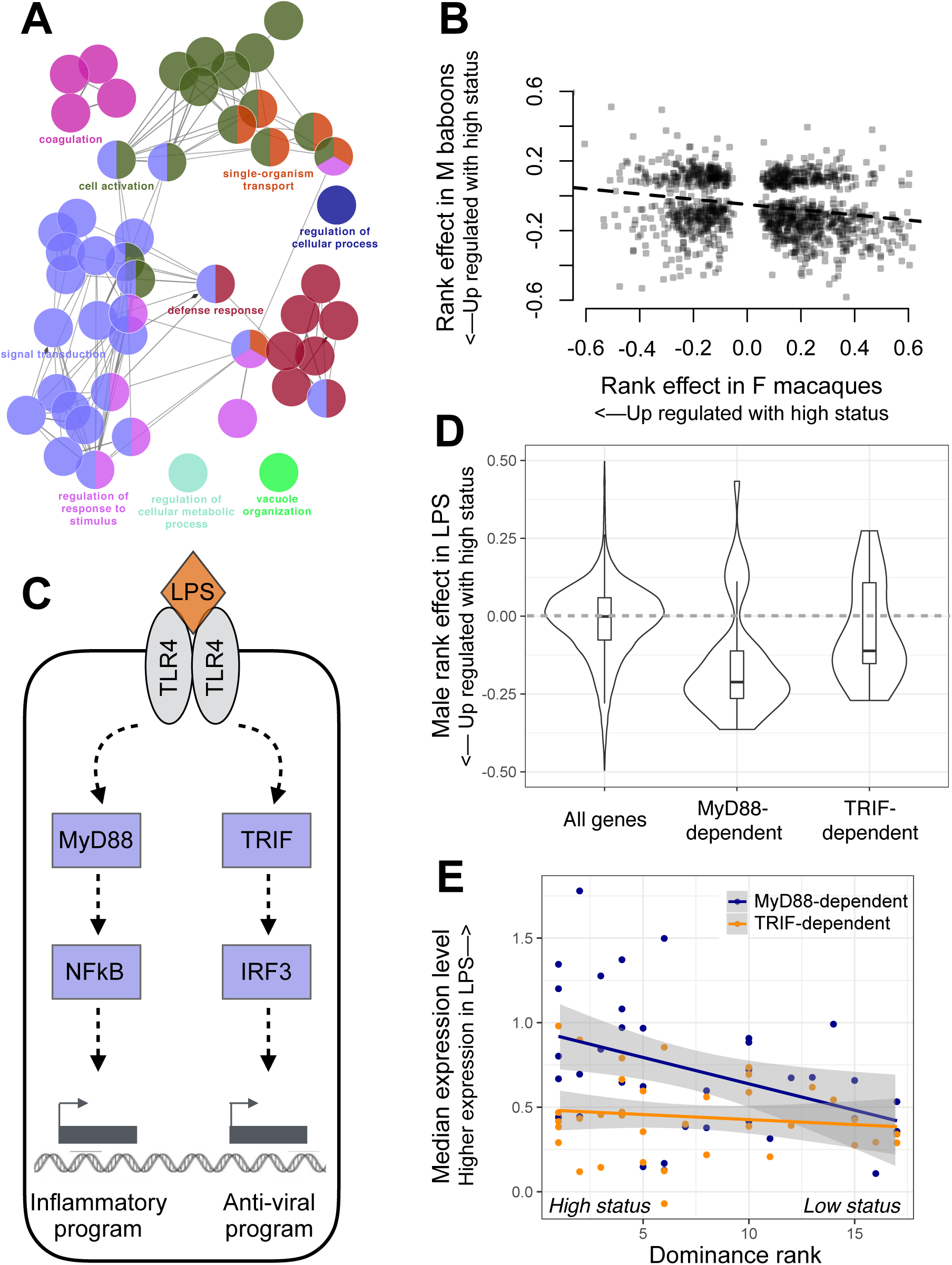
High social status in males is associated with up-regulation of innate immune genes. (A) Gene ontology (GO) term enrichment for genes that were up-regulated in high status males relative to low status males (see also Figure S4 and Tables S4-5). Inclusion is conditional on significant up-regulation by LPS. (B) X-axis: effect of rank on gene expression previously reported for captive female rhesus macaques (1), for leukocytes incubated in the presence of LPS. Y-axis: parallel results from wild male baboons. Effect sizes are plotted for genes that were found to be significantly rank-associated in both data sets and sign-reversed for the macaque data set for easier comparison to the baboon data set. (C) Simplified schematic of the LPS-induced TLR4 signaling pathway. LPS can activate a MyD88-dependent or TRIF-dependent response, leading to downstream gene regulatory responses coordinated by NFkB, IRF3, and other transcription factors. (D) Rank-associated genes that are up-regulated in the LPS condition via the MyD88 pathway (“MyD88-dependent”) are biased towards higher expression in high-status males. In contrast, the background set of all rank-associated genes (Wilcoxon rank sum test: p=5.36×10^−12^), as well as the set of rank-associated, TRIF-dependent genes (p=5.34×10^−3^), show no directional bias. (E) Male dominance rank predicts median gene expression levels across MyD88-dependent genes (rho=-0.414, p=0.012), but not across TRIF-dependent genes (rho=-0.133, p=0. 441). Each dot represents the median normalized expression level (LPS condition) for a male in the data set, plotted separately for MyD88-dependent (blue) and TRIF-dependent genes (orange).

These results were surprising given that work in captive female macaques (14, 15) and humans (39, 40) has consistently associated low, not high, social status with pro-inflammatory gene expression. Indeed, when we compared our estimates of rank effects on gene expression levels with those from captive female macaques collected using a similar method (14), genes that were more highly expressed in high status male baboons tended to be more highly expressed in low status female macaques (LPS: rho=0.222, p=2.98×10^−15^, n=1224 genes significantly associated with rank in both datasets; NULL: rho=-0.244, p=1.80×10^−7^, n=892 genes; Figure 3 and Figure S8). Further, the number of genes that displayed directionally opposite rank effects in the two data sets was significantly greater than chance expectations (binomial test, LPS: p=1.196×10^−8^, NULL: p=8.11×10^−7^).

We also observed reversal of social status effects for genes specifically involved in signaling through TLR4, the cell surface receptor for LPS. When activated, TLR4 signals through two alternative pathways: a MyD88-dependent pro-inflammatory pathway that drives an NFkB-associated transcriptional program, or a TRIF-dependent pathway that drives type I interferon regulatory activity (41). In captive rhesus macaque females, genes involved in the MyD88-dependent response are up-regulated in low status animals, consistent with social subordination-driven pro-inflammatory activity (14). In contrast, in our data set, MyD88- dependent genes tended to be more highly expressed in high status males (Fisher’s exact test, odds=1.41, p=1.22×10^−3^). Indeed, MyD88-dependent genes exhibited a stronger bias towards higher expression in high-ranking animals than either genes involved in the TRIF-dependent antigen response (Wilcoxon rank sum test: p=5.34×10^−3^) or the full set of genes analyzed (p=5.36×10^−12^: Figure 3). As a result, male dominance rank strongly predicted median gene expression levels across all rank-associated MyD88-dependent genes (rho=-0.414, p=0.012; no rank-related pattern is observable for TRIF-dependent genes: rho=-0.133, p=0. 441; Figure 3).

### Mendelian randomization analysis indicates that the gene expression signature of high status precedes attainment of high rank

Our results reveal a widespread signature of male social status in leukocyte gene expression. In contrast to experimental studies in captive animal models, however, the direction of the causal arrow connecting social status to gene expression here is unclear (18, 19, 35, 36, 42, 43). In natural populations, correlates of dominance rank can arise because being high (or low) ranking causally alters an animal’s physiology. Alternatively, in competitive ability-based hierarchies, hormonal, immunological, or other physiological correlates of rank could be indicators of condition or quality, which in turn are causal to an animal’s ability to achieve high rank (analogous to “health selection” explanations for social gradients in humans). In this scenario, up-regulation of innate immune defense genes and an increased ability to fight off infection could be an indicator of male quality that is causal to attainment of high status (19, 35).

To differentiate between these hypotheses, we took advantage of the extensive behavioral data available for our study subjects and the ability to use genotype as instrumental variables in Mendelian randomization analysis (37). We reasoned that, if social status-associated gene expression profiles are a *consequence* of male rank, rank effects on gene expression should be mediated by rank-associated behaviors (as they are in experimental studies of rhesus macaques (14)). In particular, high-ranking males in our sample expend more energy in mate guarding and physical competition, and thus initiate more agonistic behavior towards other adult males (rho=- 0.403, p=0.012; Figure 4). In contrast, low-ranking males are more often targets of agonistic behavior from other adult males (rho=0.704, p=8.15×10^−7^; Figure 4), increasing their exposure to social subordination-induced stress (36, 44). However, we found no evidence that rates of either received or initiated agonisms with other adult males explained the relationship between male dominance rank and gene expression (Figure 4). Of the 2277 male rank-associated genes tested, only 0.395 % and 0.263% were significantly mediated by initiated or received harassment, respectively, and rank effects were very similar whether we excluded or included these candidate mediators (agonisms initiated: R^2^=0.649, p<10^−10^; agonisms received: R^2^=0.916, p<10^−10^). These results suggest that rank-driven differences in agonistic behavior do not cause rank-associated variation in immune gene expression.

**Figure 4.**
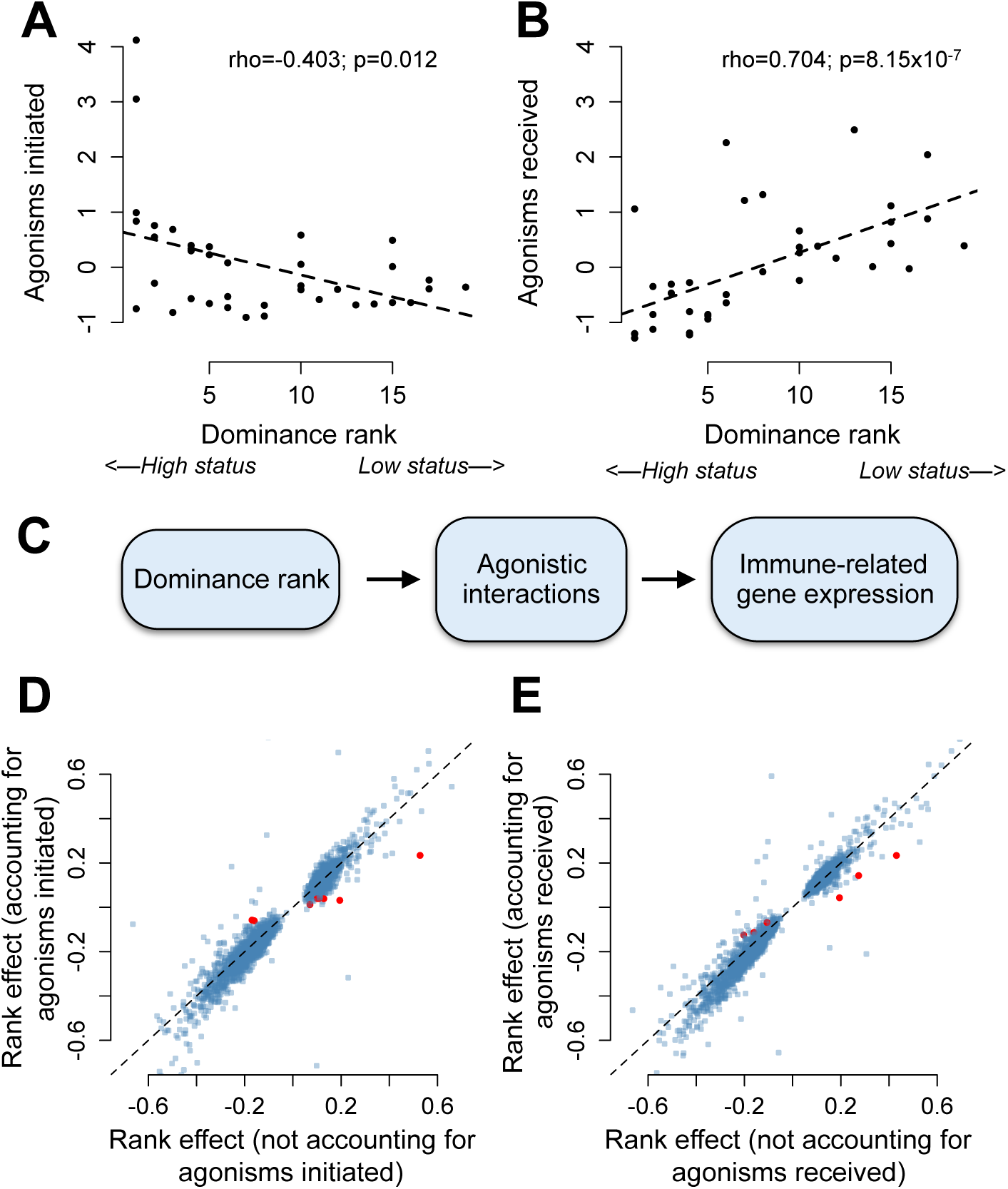
Social status-gene expression associations in males are not explained by dominance rank-associated agonistic behavior. (A) High-ranking animals in our dataset initiate more agonisms than low ranking animals (Spearman’s rank correlation, rho=-0.403, p=0.012), and (B) low-ranking animals receive more agonisms (rho=0.704, p=8.15×10^−7^). Y-axis represents a normalized measure of agonism rate corrected for observer effort. We tested the hypothesis presented in (C). However, we found that neither (D) agonisms initiated nor (E) received explain the relationship between male dominance rank and gene expression. Each point represents the estimate of the rank effect in the presence (x-axis) or absence (y-axis) of the putative mediating variable (rho=0.953 and 0.964 in panels C and D, respectively; both p<10^−10^). Of the 2277 rank-associated genes tested, only 0.26% and 0.39% (shown as red dots) were significantly mediated by initiated and received harassment, respectively. Dashed line shows the x=y line.

Alternatively, if high status-associated gene expression signatures *precede* attainment of high rank (i.e., are characteristic of high-quality males in good condition), males with genotypes that predispose them towards high expression of innate immune genes should also tend to be high ranking. If so, genetic variation should predict both gene expression and, via a path through gene expression, male dominance rank; Figure 5). This prediction can be tested using Mendelian randomization (MR), a form of instrumental variable analysis. An instrumental variable is a variable that randomly distributes an intermediate variable across study subjects, and therefore mimics case/control assignment in a randomized clinical study. In MR, the instrumental variable is always genotype, and the intermediate variable is often a molecular trait (in our case, gene expression) that is both strongly predicted by genetic variation and is hypothesized to be causal to the outcome variable of interest (in our case, dominance rank). (37, 45). For our analyses, we used projections onto the second principal component of gene expression in males as the intermediate variable, because this composite measure captures much of the dominance rank effect on gene expression (Figure 1; rho=0.44, p=1.26×10^−4^). Further, genes that load heavily on PC2 are highly enriched for genes involved in GO categories such as ‘positive regulation of inflammatory response’, ‘TLR4 signaling pathway’, and ‘defense response to Gram-positive bacterium’ (Table S7 and Figure 5). These observations indicate that PC2 can be treated as an intermediate variable for MR analysis that captures not only rank-associated variation in gene expression, but specifically rank-associated variation in innate immune defense genes.

**Figure 5.**
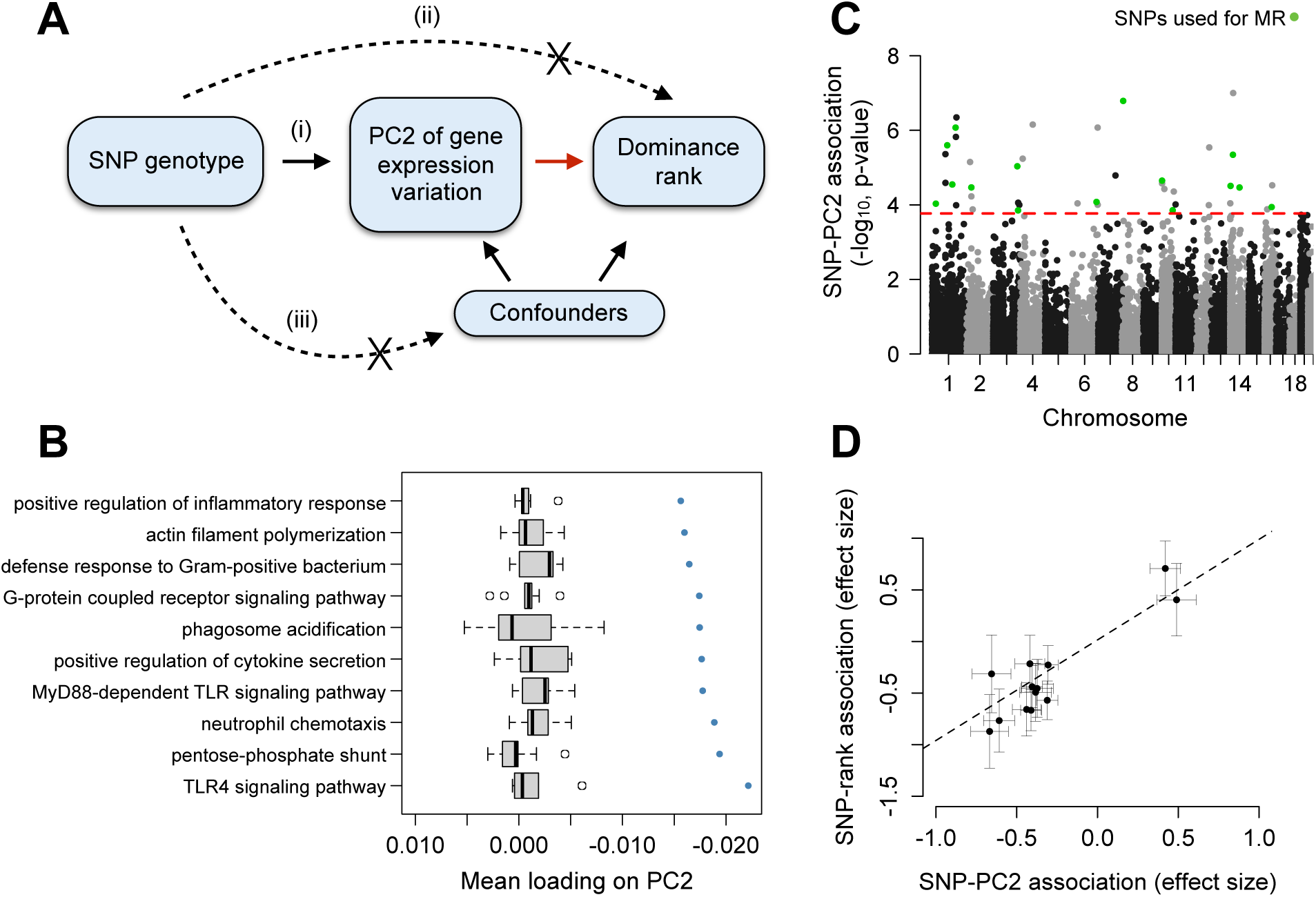
Changes in gene expression precede social status attainment in male baboons. (A) Mendelian randomization can be used to test the hypothesis, highlighted with a red arrow, that PC2 of gene expression variation (treated here as a measure of quality or condition) is causal to dominance rank. This test is robust provided that the following assumptions are met: (i) SNP genotype (the instrument) is a strong predictor of gene expression PC2, (ii) SNP genotype has no association with dominance rank except through its effects on gene expression, and (iii) SNP genotype is not related to confounding factors. (B) and (C) show that assumption (i) is satisfied. We see no detectable SNP-rank relationship for these variants independent of PC2, satisfying assumption (ii). (B) Genes that load highly on PC2 tend to be involved in innate immune signaling pathways. X-axis represents the mean PC2 loading for genes in the GO category represented on the y-axis. Blue points = observed data; grey boxplots = mean loadings for each GO category after randomizing genes across categories (while maintaining the same number of genes in each category). (C) Manhattan plot showing the strength of the association (-logio p- value) between SNP genotype and PC2 for all SNPs tested (n=29,212 candidate SNPs). Green points = SNPs that passed all filtering criteria and were used as instruments in MR analyses. Red line = genome-wide significance cutoff (5% FDR) (2). (D) Intuitively, if gene expression is causal to dominance rank, individuals with genotypes that predispose them toward low PC2 gene expression values should tend to also be high ranking (low PC2 values are associated with high social status; F************igure 1). Consistent 'with this idea, effect sizes for the SNP-PC2 relationship consistently predict effect sizes for the SNP -dominance rank relationship (MR Egger method: beta=1.2284, p=454×10^−3^).

Using the MR framework, we compared baboon males genetically randomized into a high expression class to males genetically randomized into a low expression class to evaluate the effect of immune gene expression (captured by PC2) on male dominance rank. Importantly, this approach does not imply a causal relationship between genotype and dominance rank itself; indeed, the lack of such a relationship is a requirement for MR (Figure 5; see SI Materials and Methods). To implement MR, we first identified 99,760 single nucleotide polymorphisms (SNPs) that segregated in our study population, 29,212 of which had a minor allele frequency (MAF)>5% and were not in strong linkage disequilibrium (r^2^>0.5) with other nearby (<10 kb) candidate SNPs. A subset of these variants (n=20 SNPs) satisfied stringent criteria as valid instruments for MR analysis (see Methods; Figure S9).

Genotype values at all 20 SNPs were strongly associated with PC2 (FDR<5%) and, via their effects on PC2, with dominance rank (mean PVE for the correlation between a given SNP and PC2 (± SD) = 27.28 ± 6.64%). Specifically, genotypes that predisposed individuals toward low PC2 gene expression values (which is associated with high social status; Figure 1), also consistently predicted high dominance rank (MR Egger method: beta=1.2284, p=454×10^−3^; Figure 5). Genotype at these SNPs does not explain variation in dominance rank independently of PC2 (all p>0.05; see Methods) and cannot be reverse-causally altered by dominance rank or gene expression. Thus, our analysis supports the hypothesis that the immune gene expression signature of rank precedes rank attainment. Importantly, our results are robust to the effects of outlier instruments and the potential confounding effects of genetic admixture, population genetic structure, and body size (SI Materials and Methods). Further, when we applied MR analysis to data from captive female macaques (14), in which social status is experimentally imposed, we found no support for status-associated gene expression differences as a precursor to rank, as expected (Figure S10, SI Materials and Methods).

## DISCUSSION

An increasingly large body of research shows that social status is reflected in patterns of gene regulation (10, 11, 14, 46). Our results reinforce this observation by revealing, for the first time, a strong link between dominance rank and gene expression in a natural mammal population. Despite substantial interest in the relative contribution of genetic, environmental, and demographic effects to gene expression (47), social environmental variation is rarely considered. Our results combine with those of previous studies to suggest that its omission ignores a major source of variance—one that is perhaps unsurprising in retrospect, given the centrality of social relationships to fitness outcomes in social species (2, 4).

However, our findings also stress that the social status-gene expression relationship is highly context-dependent. Previous work has emphasized the association between low social status and increased expression of innate immune and inflammation-related genes (16), but in our analyses, we observed no strong associations in females and a reversal of this pattern in males. This heterogeneity—between species, captive versus wild study systems, and sexes— points to a more nuanced relationship between social status and gene regulation than previously appreciated. Instead, it paints a picture consistent with decades of work on rank associations with other physiological outcomes, especially glucocorticoid levels. These studies emphasize that, because both the predictors and social implications of dominance rank vary across species, populations, and social contexts, so too will its physiological and fitness correlates (3, 18, 48).

One potential explanation for why we identified few rank-associated genes in female baboons, despite clear rank effects on gene expression in captive female rhesus macaques (14), involves differences in the opportunity for social support. Low social status has been most strongly linked to pro-inflammatory gene expression signatures in societies where low status individuals experience chronic social stress and elevated glucocorticoid levels (10, 16). This situation typically occurs when rank hierarchies are aggressively enforced, and low-ranking animals have little social support (3, 18, 42). The first condition (strict hierarchy enforcement) holds for both captive rhesus macaque and wild baboon females, but the second condition (absence of social support, which for females of both species is usually derived from kin), differs. In studies designed to test the causal effects of social status, captive rhesus macaque females are typically housed in groups without close kin, which breaks up the correlation between relatedness and dominance rank (49). In contrast, our study subjects were sampled from natural social groups in which females typically co-reside with their maternal relatives. Thus, low-ranking baboon females often have the opportunity to buffer themselves against status-associated stressors by investing in social bonds with kin, while captive macaque females do not. Indeed, in the same baboon population studied here, rank does not predict female lifespan independently of its effect on social integration (50). These observations suggest the importance of future studies on the molecular signatures of social integration in females, including the potential for social status-social integration interactions.

In contrast to females, we observed a strong signature of dominance rank in samples collected from male baboons. However, we found that high status males tended to upregulate inflammation and immune defense-related genes, contrary to findings in humans and captive female macaques that have reported the opposite pattern (11, 14). Further, instead of social status driving variation in gene expression, our MR analysis suggests that males who compete successfully for high rank are already non-random with respect to immune gene expression. Several explanations could account for this result. First, the ability to maintain an energetically costly pro-inflammatory state could be an indicator of male condition. In Amboseli, high ranking male baboons exhibit high testosterone levels, and alpha males also exhibit high glucocorticoid levels (36). Given that glucocorticoids, and perhaps testosterone, can suppress some immune responses, males that can maintain active innate immune defense systems in the face of these hormone signals may be particularly high quality (51, 52). Notably, male baboons in Amboseli also exhibit graded increases in glucocorticoid levels with decreasing social status and some evidence for glucocorticoid resistance (36, 44). Our observation that inflammation-related gene expression is nevertheless higher in high-ranking males thus suggests that glucocorticoid and immunological correlates of rank can be decoupled, perhaps as a function of male condition.

A non-mutually exclusive explanation is that individuals that mount strong innate immune defenses are more likely to succeed in, and recover from, physical conflicts with conspecifics. Indeed, at any given time after injury, the highest ranking males in Amboseli are three times more likely to heal from wounds and injuries than low ranking males (35). Upregulation of genes involved in the inflammatory response, which facilitates pathogen defense and recovery from tissue damage, may explain this observation. Males who can maintain high expression of inflammation-related genes may therefore enjoy an adaptive advantage in the fights required to attain and maintain high status. Such an explanation does not exclude the possibility that a persistent pro-inflammatory state is also costly, consistent with its role in aging and susceptibility to disease (17). However, because dominance rank is the best predictor of reproductive success in male baboons (23), even physiological states that incur long-term costs will be selectively advantageous if they aid in competition for high rank. Previous hypotheses have predicted higher investment in inflammatory defenses for low ranking males than high ranking males (53, 54), an argument that has received mixed support in empirical studies (19). Our results directly oppose this idea and suggest that the relationship between inflammation and social status is highly context-dependent.

Finally, our study highlights a simple functional genomic approach for investigating how the social environment and other individual characteristics covary with immune function in natural populations. *Ex vivo* challenge experiments like the one conducted here can be used to measure physiological sensitivity to predictors of interest, test hypotheses about trade-offs in investment, and identify candidate molecular mechanisms that link the environment to health or fitness-related outcomes. In addition, because gene regulatory phenotypes are far more amenable to trait mapping approaches than organism-level traits, they can be integrated with Mendelian randomization to investigate the causal direction linking gene regulatory phenotypes to other traits. This approach is increasingly used in humans when experimental randomization is not possible (37, 45), but has not been applied in observational studies of other species, which are often confronted with the same limitations.

## METHODS

We obtained blood samples from 61 adult baboons using previously described procedures (55). All animals were individually recognized members of a long-term study population that has been monitored by the Amboseli Baboon Research Project (ABRP) for almost five decades (34). From each animal, we collected: (i) whole blood used to isolate peripheral blood mononuclear cells (PBMCs) for flow cytometry analysis and (ii) 1 mL of whole blood in each of two TruCulture tubes (LPS or NULL), which were subsequently incubated for 10 hours at 37°C and used to isolate leukocytes (for mRNA-seq) and serum (for cytokine profiling). Dominance hierarchies were constructed monthly for every social group in the study population based on the outcomes of dyadic aggressive encounters. Ordinal dominance ranks were assigned to every adult based on these hierarchies, such that low numbers signify high rank/social status and high numbers signify low rank/social status (28).

mRNA-seq data were mapped to the anubis baboon genome *(Panu* 2.0). The resulting counts were filtered to remove lowly expressed genes and normalized to remove batch and cell type composition effects. To identify social status-associated gene expression variation, we used principal components analysis to investigate rank associations with the major axes of variation in the full gene expression dataset and linear mixed-effects models to test for rank associations at each gene. In the mixed models, we nested rank within sex and included age (nested within sex) and condition (NULL or LPS) as fixed effects covariates. A relatedness matrix inferred from SNP genotypes was included to control for genetic relatedness among our study subjects.

To ask whether status-associated behaviors in males explained observed relationships between dominance rank and gene expression, we used long-term behavioral data on agonism rates in formal mediation analyses. Specifically, we compared the estimate of the rank effect on each gene in a model that included versus excluded the mediator (agonisms initiated or agonisms received), and used bootstrap resampling to assess significance. To ask whether variation in gene expression precedes variation in dominance rank, we used Mendelian randomization (MR), a form of instrumental variable analysis (37). To conduct MR, we combined the pipeline and filtering criteria outlined in Figure S9 with the MR Egger method (56) implemented in the R package ‘MendelianRandomization’ (57).

All statistical analyses were performed in R (58). Further details for all experimental and statistical procedures can be found in the SI Materials and Methods. Data included in this study were obtained in accordance with Institutional Animal Care and Use Committee protocols approved by Duke University (IACUC A004-15-01).

## ACKNOWLEDGMENTS

We gratefully acknowledge the National Science Foundation and the National Institutes of Health for their support, currently through NSF IOS 1456832, NIH R01AG053330, R01HD088558, and P01AG031719. This work was also supported by the Leakey Foundation, BCS-1455808, and R21-AG049936. We thank members of the Amboseli Baboon Project, especially J. Altmann for her foundational role in establishing these data sets; R.S. Mututua, S. Sayialel, J.K. Warutere, T. Wango, and V. Oudu in Kenya; N. Learn, J. Gordon, and K. Pinc for expert database management and design; and T. Voyles and C. Del Carpio for assistance with wet lab work. Thanks to N. Snyder-Mackler, J. Anderson, X. Zhou, L. Barreiro, S. Mukherjee, and J. Sanz for support on data analysis and feedback on the manuscript. Finally, we thank the Baylor College of Medicine Human Genome Sequencing Center for access to the baboon genome assembly *(Panu 2.0)*. For a complete set of acknowledgments, please visit http://amboselibaboons.nd.edu/acknowledgements/.

## AUTHOR CONTRIBUTIONS

AJL, JT, and EAA designed research; SCA, EAA, and JT contributed long-term data; AJL, MYA, RN, PM, and FN performed research; TK contributed research/analytic tools; AJL analyzed data; and AJL, EAA, and JT wrote the paper, with contributions from all co-authors.

